# A New Classification Method Based on Dynamic Ensemble Selection and its Application to Predict Variance Patterns in HIV-1 Env

**DOI:** 10.1101/2022.01.31.478521

**Authors:** Mohammad Fili, Guiping Hu, Changze Han, Alexa Kort, John Trettin, Hillel Haim

## Abstract

Therapeutics that target the envelope glycoproteins (Envs) of human immunodeficiency virus type 1 (HIV-1) effectively reduce virus levels in patients. However, due to mutations, new Env variants are frequently generated, which may be resistant to the treatments. The appearance of such sequence variance at any Env position is seemingly random. A better understanding of the spatiotemporal patterns of variance across Env may lead to the development of new therapeutic strategies. We hypothesized that, at any time point in a patient, positions with sequence variance are clustered on the three-dimensional structure of Env. To test this hypothesis, we examined whether variance at any Env position can be predicted by the variance measured at adjacent positions. Sequences from 300 HIV-infected patients were applied to a new algorithm we developed. The k-best classifiers (KBC) method is a dynamic ensemble selection technique that identifies the best classifier(s) within the neighborhood of a new observation. It applies bootstrap resampling to generate out-of-bag samples that are used with the resampled set to evaluate each classifier. For many positions of Env, primarily in the CD4-binding site, KBC accurately predicted variance based on the variance at their adjacent positions. KBC improved performance compared to the initial learners, static ensemble, and other baseline models. KBC also outperformed other algorithms for predicting variance at multi-position footprints of therapeutics on Env. These understandings can be applied to refine models that predict future changes in HIV-1 Env. More generally, we propose KBC as a new high-performance dynamic ensemble selection technique.

## 1. INTRODUCTION

Four decades after recognizing HIV-1 as the causative agent of acquired immune deficiency syndrome (AIDS), this virus is still a major health concern worldwide. In the year 2019, 38 million individuals were living with HIV, 690,000 died from AIDS-related disease, and 1.7 million were newly infected [1]. To treat HIV-infected individuals, multiple therapeutics are available; they bind to HIV-1 proteins and can effectively inhibit their function. However, the replication machinery of HIV-1 is highly prone to errors. As a result, new variants of its proteins are generated, some of which contain changes at the sites targeted by the therapeutics [2]. Subsequent expansion under the selective pressure of the therapeutic can lead to clinical resistance [3, 4]. Since the appearance of the mutations is random, the emergence of resistance by changes at any position of an HIV-1 protein is considered unpredictable. There is a critical need to better understand the patterns of change in HIV-1 within the host. Such knowledge can lead to the design of new strategies that tailor treatments to infected individuals based on the properties of the infecting virus and the changes expected to occur. Multiple tools have been developed over the past two decades to predict the evolution of other viruses, primarily influenza virus; they aim to forecast changes in structural properties of virus proteins that can inform the design of vaccines [5]. Unfortunately, the number of tools developed to model and predict the changes in HIV-1, and particularly within the host, is limited [6–9].

### 1.1. Toward a Better Understanding of the Variance Patterns in HIV-1 Env Within the Infected Host

Of all HIV-1 proteins, the envelope glycoproteins (Envs) exhibit the highest level of diversity, both within and between hosts [10, 11]. Env adorns the surface of HIV-1 particles and allows the virus to enter cells [12]; it is thus a primary target in AIDS vaccine design [13]. Env is composed of approximately 850 amino acids (some diversity in length exists between different strains). In the infected host, new amino acid variants continuously appear at multiple positions of this protein. Consequently, at any time point during chronic infection, 10% or more of Env positions can exhibit variance in amino acid sequence between co-circulating strains [14, 15]. The random nature of the mutations, the extreme diversity of Env within and between hosts, and the structural complexity of this protein limit our ability to model the changes.

Whereas the amino acid sequence at any Env position can vary between strains in different hosts, the level of in-host variance in amino acid sequence at each Env position is similar in different individuals [9]. Variance at each Env position is specific for each subtype (clade) of HIV-1. Thus, patterns of variance in the host are not merely random “noise” but reflect the inherent properties of the virus. Variance describes the permissiveness of sites to contain amino acids with different chemical properties, which reflects the strength of the selective pressures applied to them. Such pressures are typically not applied on individual positions but on multi-position domains of Env. In this work, we investigate the spatial clustering of variance across the Env protein. Specifically, we test the hypothesis that the absence or presence of sequence variance at any position of Env can be predicted based on the variance at adjacent positions on the three-dimensional structure of the protein. If the tendency for co-variance of adjacent positions is stable over time, then the patterns of variance detected in a patient may provide insight into future changes. To test the above hypothesis, we developed a new algorithm that selects the best classifiers for a new observation using a dynamic mechanism.

### 1.2. Multiple Classifier Selection Algorithms

As the complexity of any dataset increases, the ability of any single classifier to capture all patterns is reduced. Therefore, it is necessary to integrate multiple classifiers for improved classification accuracy. However, the use of the same set of classifiers statically over the whole feature space can affect the overall performance of an algorithm. In fact, a classifier may perform well in some subspaces of the data but exhibit poor performance in others. One solution to this problem is the selection of the best classifiers out of a pool of existing learners dynamically based on some competitiveness criteria and to use this subset of classifiers for predicting the class label of a new observation in a region. Such a mechanism will help to select the best classifiers in each subspace.

A considerable amount of work has been conducted in the field of multiple classifier systems (MCS) during the past two decades. MCSs are divided into two basic categories: static and dynamic. Static classification techniques use the same classifiers over the whole range of the dataset, while dynamic classification methods use the best classifier(s) at local regions. The method we propose in this work is built upon the dynamic classification approach [16, 17]. Dynamic classification can be further divided into dynamic classifier selection (DCS) and dynamic ensemble selection (DES). The DCS approach uses one base classifier to predict class labels in a local region [18], while the DES methods use many base classifiers [19]. Here, we propose a DES-based classification technique.

DES methods are composed of three main steps: (i) classifier generation, (ii) ensemble selection, and (iii) classifier combination. For the first step, either heterogenous classifiers [20–22] or homogenous ones [23–25] can be used. The proposed method will use decision trees as the homogenous classifiers for its classifier generation phase. Since local accuracy is a key feature of the DES method, many algorithms use k-nearest neighbors as a framework [19, 26, 27]. Other methods for generating different homogeneous classifiers have been proposed, including random subspace [28], bagging [29], boosting [30], and clustering [31].

The second step in DES design is ensemble selection. Some selection mechanisms utilize probabilistic models that adopt the probability of correct classification as the measure of competence [32]. Alternatively, one can use a wide range of classifiers to increase diversity, which increases accuracy as a result [33]. Models based on accuracy are another type of selection method that considers the classifiers with the highest levels of accuracy in the local area [17].

The third step of a DES method is classifier combination, which aggregates the gathered information into a single class label. Dynamic classifier weighting is one aggregation technique [34, 35]. Artificial neural networks have also been adopted to aggregate the results of base learners and use a cost function to maximize accuracy [36]. The majority vote method has also been used, which applies predictions of the highest accuracy classifiers in a local region and then takes the majority of these predictions to determine its output [37].

In this study, we introduce a novel approach to use a dynamic mechanism to increase the flexibility of the algorithm. The best classifiers are chosen based on the performance of the initial learners in the neighborhood of a new observation. Bootstrapping has been adopted to increase randomness, thus introducing more diversity within the base learners. We also introduced a classifier scoring approach, upon which the selection decision of a classifier can be made. To define a neighborhood for a new observation, we used the k-nearest neighbors (KNN) algorithm. Each observation has a feature vector that is used for the neighborhood selection process. The novelty of this method is in the dynamic classifier selection approach, where we introduced a weighting mechanism to evaluate each classifier’s performance within a neighborhood of an instance and decide if the classifier is good enough to be used for prediction. This approach is based on bootstrap resampling, which creates the out-of-bag sample that can be used along with the resampled data in the classifiers’ evaluation process.

We tested the KBC algorithm with a panel of sequences from HIV-1 infected individuals. These data describe for each patient the absence or presence of variance at each position on the three-dimensional structure of the Env protein. We examined whether the variance at each position (or at a group of positions) can be predicted based on variance at adjacent positions on the three-dimensional structure of the protein. In many cases, the KBC method showed improvement in the classification metrics relative to other machine learning algorithms. Considerably higher performance was observed in the CD4-binding domain of Env, which is the target of multiple antibody therapeutics against HIV-1 [38–42]. The primary advantage of KBC is that it does not use the same set of classifiers for the entire problem space; instead, it identifies the subset of learners that are capable of better predicting class labels for the observations in a region. This flexibility helps to avoid the loss of helpful learners and to limit the retention of weak learners that occurs when a fixed number of classifiers is applied.

## 2. METHODS

### 2.1. Datasets

In this study, we developed a novel dynamic classification approach to predict the variance at any Env position based on variance at adjacent positions on the protein. Because of the folded structure of the protein, the physical distance between any two positions (in Ångstroms, Å) is used as the measure of proximity. These distances are based on the cryo-electron microscopy (cryo-EM) coordinates of the Env protein. A description of the datasets is provided in the following sections.

#### 2.1.1. HIV-1 Env Sequence Data

Nucleotide sequences of the HIV-1 *env* gene were obtained from the National Center for Biotechnology Information (NCBI) database (https://www.ncbi.nlm.nih.gov) and from the Los Alamos National Lab (LANL) database (https://www.hiv.lanl.gov). Sequence data for HIV-1 clades B and C were downloaded and processed separately. The clade C dataset is composed of 1,960 sequences from 109 distinct patients. The clade B dataset is composed of 4,174 sequences from 191 distinct patients. For each patient sample, all Envs isolated were analyzed (6-30 sequences per sample). All *env* genes were cloned from the samples by the single genome amplification approach [43] and sequenced by the Sanger method. Sequences of non-functional Envs were removed, as were all sequences with nucleotide ambiguities or large deletions in conserved regions [9, 44]. Nucleotide sequences were aligned using a Hidden Markov Model with the HMMER3 software [45] and then translated into the amino acid sequence, which was used for the analysis. All 856 Env positions described in the manuscript conform to the standard HXBc2 numbering of the Env protein [46]. Potential N-linked glycosylation sites (PNGSs) contain the sequence motif Asn-X-Ser/Thr, where X is any amino acid except Pro. To account for the presence of N-linked glycans on the Asn residues, the first position of all Asn-X-Ser/Thr triplets was assigned a unique identifier. All aligned sequences from each patient were compared to determine whether each of the 856 positions has variance in amino acid sequence (position is assigned a variance value of 1) or whether all sequences from that patient sample have the same amino acid at the position (assigned a variance value of 0).

#### 2.1.2. Env Structural data

The Env protein is folded so that positions from different domains can be close in the three-dimensional structure of the protein. To determine the positions closest to each position of interest, we used the coordinates of the cryo-EM structure of Env. For clade B viruses, we used the coordinates of Env from HIV-1 clade B strain JRFL (Protein Data Bank, PDB ID 5FUU) [47]. For clade C viruses, we used the coordinates of Env from HIV-1 clade C strain 426c (PDB ID 6MZJ) [48]. The distance between any two positions was measured using the coordinates of the closest two atoms of the two amino acids. These data were applied to identify the 10 closest positions to each position of interest.

### 2.2. The KBC method

In the KBC algorithm, available information is utilized regarding the base classifiers for the data points in the neighborhood of a new observation to predict the class labels. This allows us to identify the classifiers that perform well in the neighborhood of a new observation and apply only those learners for the prediction. Classifiers do not perform the same in different regions of the problem space, especially when the data structure is complicated. This becomes even more tangible when more naïve learners are used. KBC applies the information from the training phase to identify those classifiers that are more helpful in finding the class label of a new instance based on their performance within a specific neighborhood. The foundation of this method, like other dynamic ensemble selection techniques, relies on three main steps: classifier generation, selection, and aggregation.

#### 2.2.1. Classifier Generation

In the proposed method, we used the decision tree algorithm as the base learner. We selected this algorithm primarily because of its training speed. In general, for the first step, *M* base learners, *L*_1_, *L*_2_,…, *L_M_* are generated. The number of base learners is a hyperparameter of the KBC model.

In the next step, to identify the best classifiers, we used bootstrap resampling. In this manner, we can reserve an untouched sample (i.e., the out-of-bag sample, OOB), which is used along with the resampled observations for the selection of the best classifiers. For the training set, *X^train^*, we have *x*_1_, *x*_2_,…, *x_N_* feature vectors (one for each observation), and *y*_1_, *y*_2_,…, *y_N_* as their class labels. We denote the test set as *X^test^*.

We use bootstrapping to train each base learner *L_i_*. This approach creates two separate sets of observations for learner *L_i_*: a resampled set 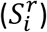 and an OOB set 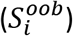. In other words, we make a partition:

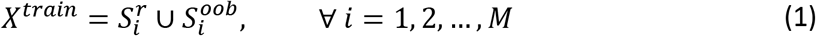

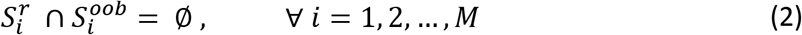

If we define an event *A*, as a data point *x_j_* (*j* = 1,2, …, *N*) that belongs to the OOB sample:

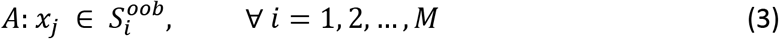

then the probability of such event can be calculated as:

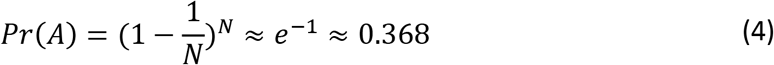

where, *N* is the total number of observations in the training set *X^train^*. Later, we will show how to use this information in the algorithm as a starting point.

To introduce more randomness among the base learners, we used random sampling for feature selection. To accomplish this task, the algorithm randomly picks *f* features out of all available attributes for *L_i_*. In other words, the learner *L_i_* is trained over the subset of the features of 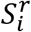 which is denoted by 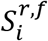. Knowing the set of *f* features for the learner *L_i_*, one can also create 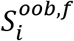 for the evaluation phase.

#### 2.2.2. Classifier Selection

First, each base learner is used to predict all instances in *X^train^*, including the resampled and OOB data. Then, the classification results are mapped onto a binary variable, *z_ij_*, which is 1 or 0 based on whether the classifier *L_i_* correctly classified the instance *x_j_* or not, respectively:

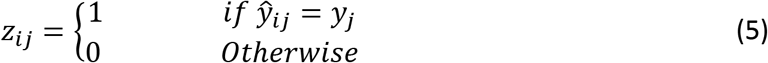

where, *i* = 1,2,.., *M* and *j* = 1,2,…, *N* and 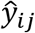 is the class label that is predicted by the learner *L_i_* for *x_j_* ∈ *X^train^*. The result of this phase is an *M*N* binary matrix *Z*, in which each row represents the mapped prediction result for one base learner, and each column corresponds to an observation in the training set, *X^train^*:

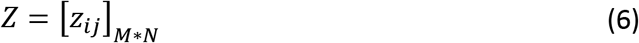

For efficiency, we perform this only once for all observations rather than during each iteration. In effect, not all observations will be used for selecting the best classifiers, but only the ones in the neighborhood of the new observation *x_q_* ∈ *X^test^*. To find the neighbors, different approaches can be applied, one of which is the KNN method. KNN finds the closest data points to the observation of interest. However, since it is based on the distance matrix, for very large datasets, the process can take much longer than a dataset with a small number of observations.

By defining 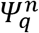 as the neighborhood of a new data point *x_q_* which includes n-closest observations, we can define:

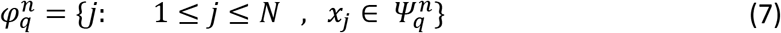

where, 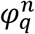 is the set of *n* indices for the data points within the neighborhood of the new instance *x_q_*.

To emphasize the contribution of correctly predicting an observation in the OOB set relative to the resampled set, we can assign greater weights to the observations in the OOB set. In this manner, we can also define the level of emphasis by tweaking the weights. However, the optimal value can be found by hyperparameter tuning.

Weighting of the OOB and resampled sets can be described by:

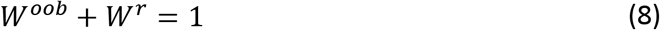

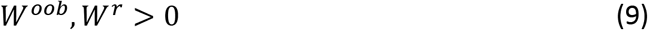

where, *W^oob^* and *W^r^* are the weights for observations within the OOB 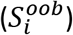 and resampled sets 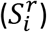 for learner *L_i_*, respectively. From **Equation 4**, we can conclude that the probability of a data point belonging to 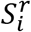 is approximately 0.632. Therefore, we can use this value as the default *W^oob^*; however, the optimal value for this parameter can be obtained via hyperparameter tuning. In general, the higher the OOB weight, the greater the focus on the OOB observations rather than the resampled set.

Now, consider the matrix *⊓* in which the type of data points (i.e., being from the OOB or resampled set) is stored:

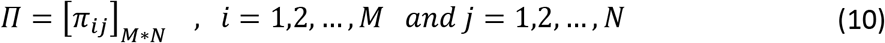

where, *π_ij_* is defined as:

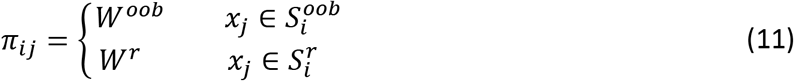

where, *i* = 1,2, …, M and *j* = 1,2, …, *N*. In the next step, the classifier’s score, *CS_i_*, can be calculated for base learner *L_i_*:

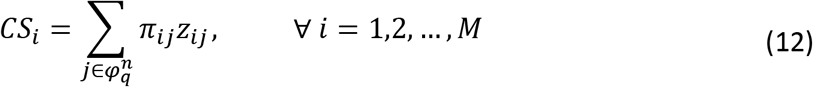

Then, we normalize the scores to the same scale:

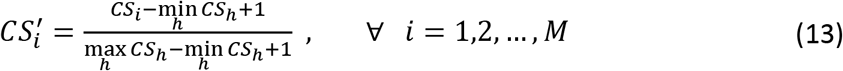

where, *h* = {1,2,…, *M*} is the set of all classifiers. This normalization rescales all scores into a range of (0, 1], and facilitates the comparison:

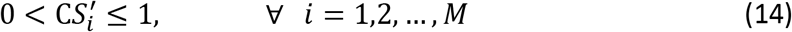

Here, 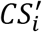 quantifies the relative importance of base learner *L_i_* to the best classifier. For instance, if all learners perform the same in a region, then we expect a value of 1 for all (i.e., no distinction between the classifiers).

**Figure 1** shows the relationship between the range of classifiers’ scores with achievable normalized scores. For instance, if the difference between the maximum and minimum classifier’s score is 1, then the normalized score of any classifier will be between 0.5 and 1. The yellow region in **Figure 1** shows all possible normalized scores for any difference between the best and worst classifiers. The blue line on the left margin of the yellow region indicates the lower bound of the normalized score for any specific normalized score’s range.

**Figure 1.**
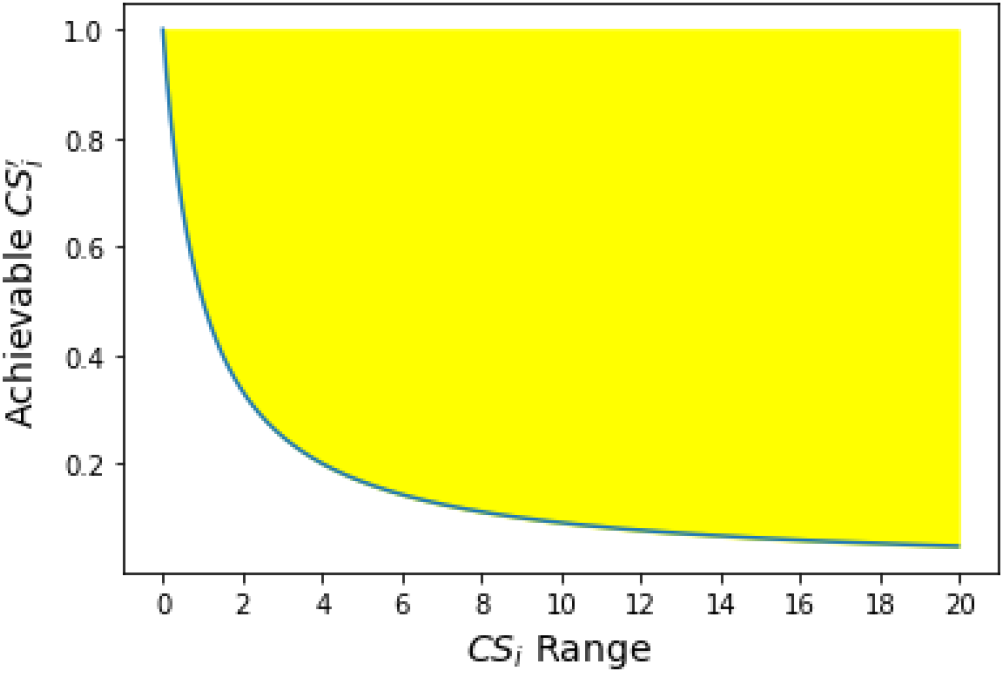
Achievable normalized scores 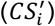 for different values in the *CS_i_* range (difference between the best and worst classifiers’ scores). The yellow region shows all possible values.

In the following equation, we can observe that, as the difference between the performance of the best and worst classifiers increases, there is greater confidence that classifiers with higher scores are performing significantly better relative to those with lower scores:

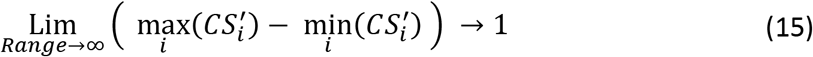

This occurs when the performance of the best and worst learners differs substantially. In such a case, we are more confident about the significance of the classifier with the high normalized score. As shown in **Equation 15**, for this extreme case, where the range approaches infinity, the limit of the difference between the best and worst normalized classifiers’ score converges to the maximum value of 1. In other words, based on **Equation 14**, we can conclude that the best and worst classifiers are at the two ends of the interval. On the other hand, if the range of scores is 0 (i.e., all classifiers have the same performance), the normalized scores will be 1 for all of them (no distinction).

Finally, based on a minimum acceptance threshold *δ*, only those classifiers with scores higher than that value will be selected. Therefore, for different observations, we expect to have a different array of scores for base learners’ performance within the neighborhoods. Thus, the algorithm will select the best classifiers based on the problem space, observations, and the base learners’ capabilities to correctly classify similar instances each time.

In **Equation 16**, the index corresponding to the k-best classifiers (out of *M* existing learners), for *x_q_*, is defined.

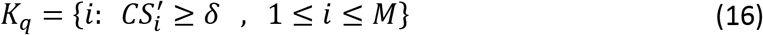

The number of best classifiers can differ from observation to observation. However, for similar points (i.e., observations within a similar segment of the problem space), we expect to have a similar set of best classifiers for prediction.

#### 2.2.3. Classifier Aggregation

Finally, once classifiers for the prediction are defined, we apply an aggregation method to obtain a single result for the new instance. Here we use the majority vote approach. For a general case in which we have *P* classes we can write:

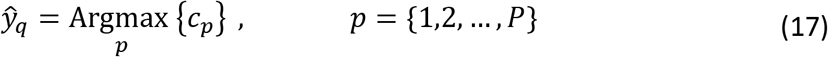

where, 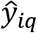 is the predicted class for the new observation *x_q_*, and *c_p_* counts the number of base learners predicting class *p*. We can write it as:

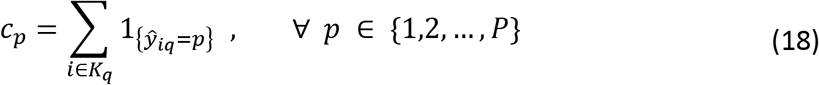

where, 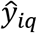 is the class label predicted by learner *L_i_* for *x_q_*.

#### 2.2.4. Hyperparameters

For the KBC algorithm, hyperparameters were designed to accommodate the variability in the dataset to ensure optimal performance. The hyperparameters of the algorithm are shown in **Table 1**. *M* is the number of base learners. If sufficient diversity exists within the base learners (i.e., among the decision trees generated), more learners typically lead to better results. We can also change the number of features (*f*) for each classifier. Using all available features can result in less randomness and reduces the level of diversity. On the other hand, using too few features, such as the extreme case of *f*=1, can result in a naïve learner that may not be much better than the random guess. However, by using a suitable fraction of the available features for training the classifiers, we can add variation between the classifiers and also increase confidence that each classifier will perform well.

**Table 1.** Hyperparameters of the KBC algorithm

| parameter | Domain | Description |
| --- | --- | --- |
| $M$ | $\in \mathbb{N}$ | Number of initial base learners |
| $f$ | $\in \mathbb{N}$ | Number of features to be selected randomly for each base learner |
| $n$ | $\in \mathbb{N}$ | Number of close neighbors to a new instance (similar instances) |
| $W^{oob}$ | $[0,1]$ | Weight of OOB instances. (Default=0.632) |
| $\delta$ | $[0,1]$ | Minimum acceptance threshold for a base learner's score to be selected in a neighborhood of a new data point. |

The third hyperparameter is the number of neighbors for a new instance (*n*). Increasing the number of neighbors to all training observations (*N*) will lead to the majority vote for a fixed set of classifiers. In this case, we expect to have no variance but high bias. At the other extreme, if we use only one neighbor, the variance will be high. Therefore, it is a bias-variance trade-off, and selecting the optimal *n* is essential for performance. The effect of this parameter on the accuracy of the model is also explored in this study.

The weight of OOB instances (*W^oob^*) plays an important role in emphasizing the unseen data for selecting the best classifiers. Since the data in the OOB set are not used during training of the base learners, predicting them correctly is more important than the observations in the resampled set. Choosing the weight as 1 will completely ignore the resampled data, whereas a weight of 0 will result in using only the resampled data in the training phase. The optimized value for the OOB weight can be obtained by hyperparameter tuning.

The last hyperparameter of the KBC model is the minimum acceptance threshold (*δ*), which determines the sensitivity in selecting the best classifiers. The higher the threshold, the smaller the number of learners we expect to have. In such a case, the variance may increase; however, at the same time, we are more confident about the set of selected classifiers in that region. On the other hand, using smaller values for *δ* may result in more learners, which reduces variance.

A summary of the KBC algorithm is shown in **Figure 2**. As explained, we start with the training set. By applying bootstrap resampling, we create OOB samples as well as a resampled set for each learner. Then we train classifiers separately on their own resampled data. In the prediction phase, for each new data point, we first identify the closest neighbors. Then, based on whether a point belongs to the OOB or resampled set, the method assigns weights to the 0-1 mapping of the initial predictions (1 for correctly classifying an observation and 0 for misclassifying it). This approach introduces a scoring function that is used for evaluating the classifiers. Finally, according to a minimum threshold acceptance value, only those classifiers for which the normalized score exceeds the selected limit will be chosen for classifying the new instance. The individual predictions are then aggregated into a single result by the majority vote aggregation method. In **Figure 3**, we show the pseudocode for the entire KBC process.

**Figure 2.**
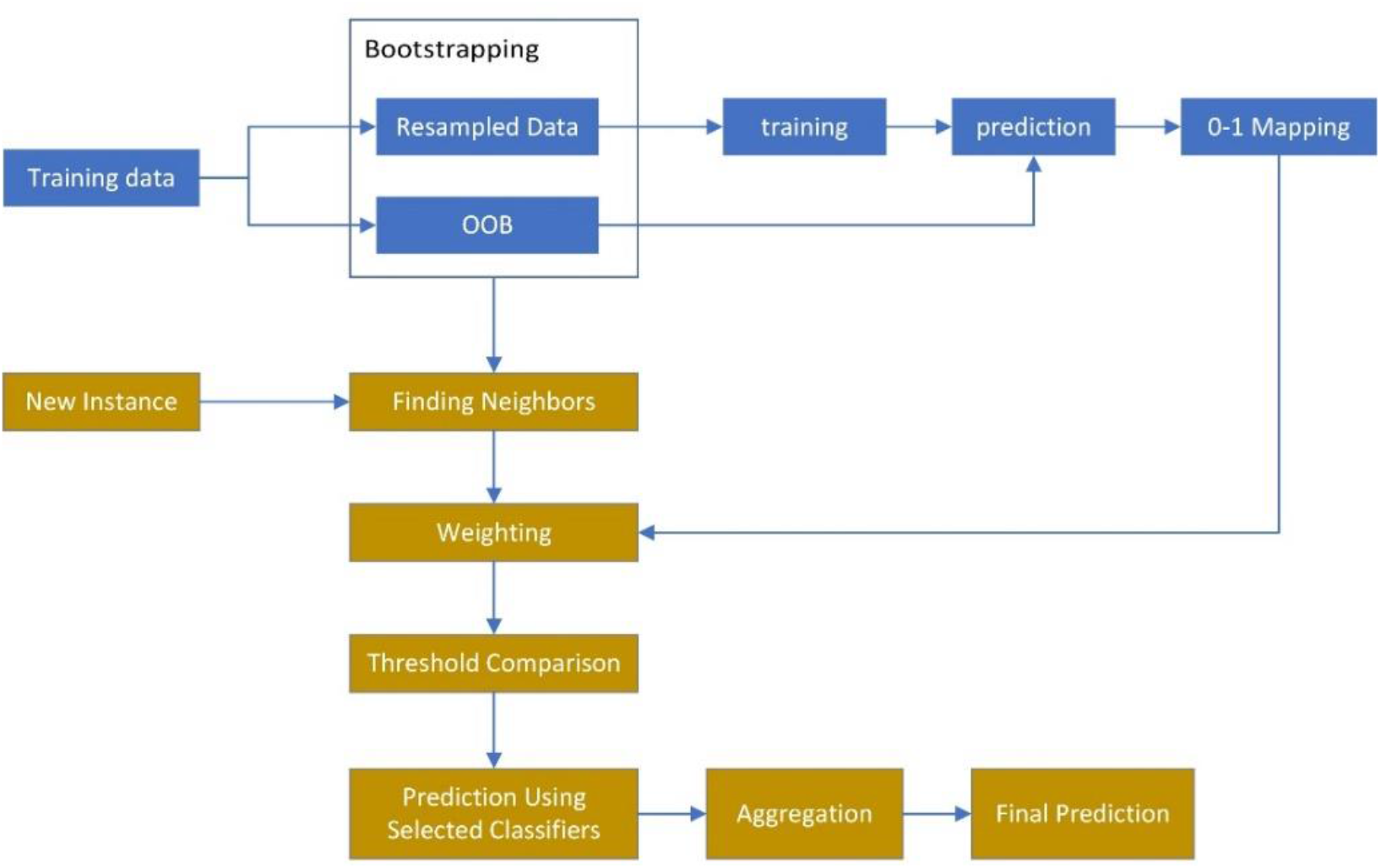
Flow chart of the KBC algorithm. It starts with the bootstrap resampling for each base learner. Then, for a neighborhood of a new data point, the weights are assigned to the 0-1 mapped values and then aggregated into a single score for each learner. Those classifiers surpassing the minimum threshold are selected for the classification of the new observation.

**Figure 3.**
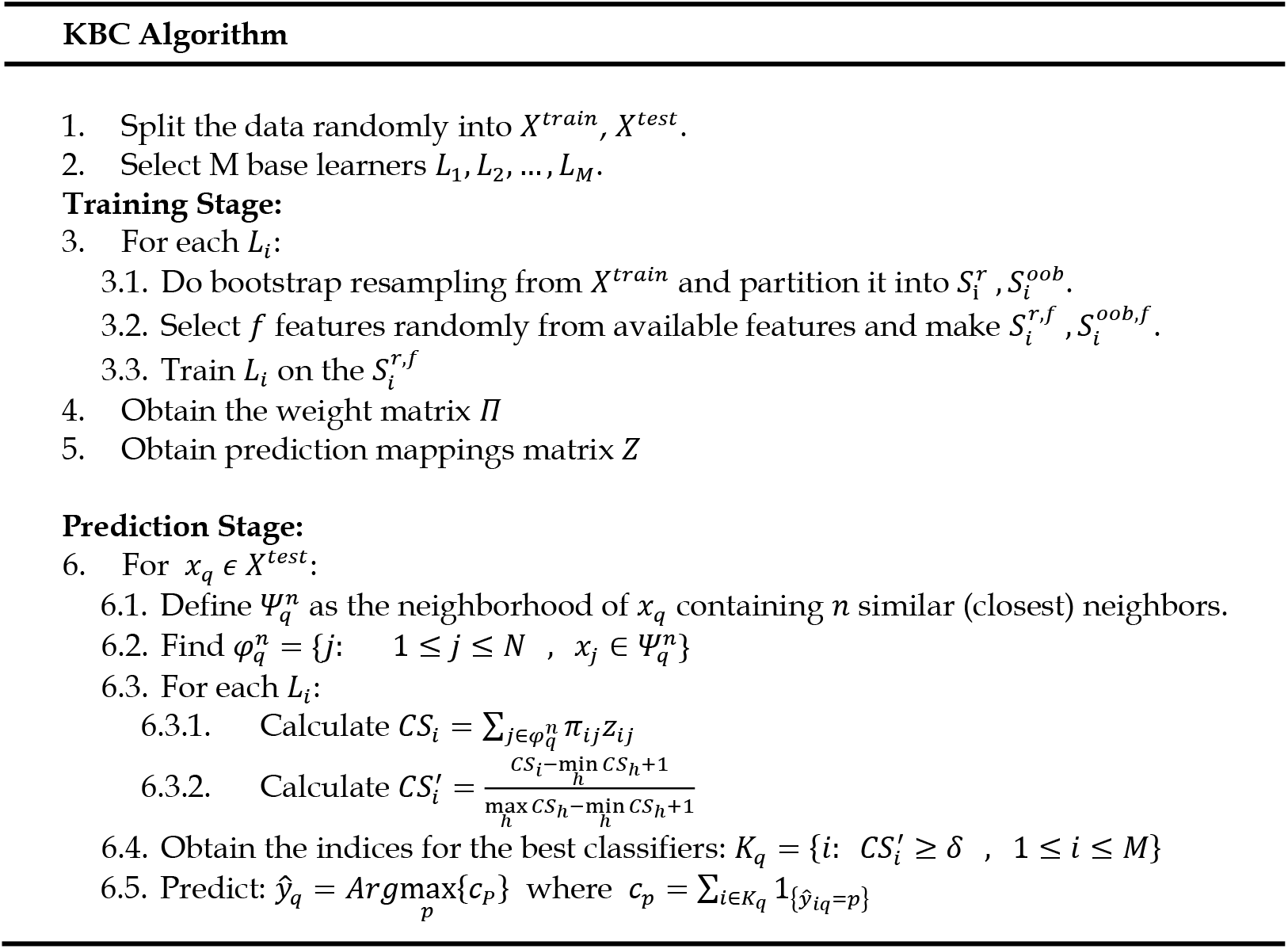
Pseudocode for the KBC algorithm.

We note that in the KBC method, one can use any set of homogenous or heterogeneous base learners. Since we used the decision tree as the base learner, we added the maximum depth of the tree to the set of existing hyperparameters and tuned the KBC model for the best performance.

### 2.3. Evaluation metrics

In this study, 5 classification metrics are used: accuracy, precision, recall, F1 score, and balanced accuracy. Accuracy depicts the percent of predictions that are correct. Precision describes the percentage of correct classifications from the group of instances that are predicted as the positive group. Recall or sensitivity represents the correct classification rate from the group of true positive instances. The F1 score takes both the precision and recall metrics into consideration. Since the HIV-1 datasets are not balanced (i.e., for any position, the proportion of variance-positive and no-variance samples is not equal), we also used balanced accuracy, which is an average of sensitivity and specificity.

The formulae for these metrics are shown:

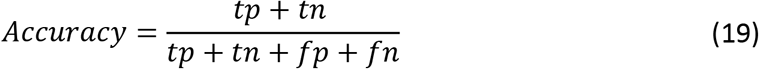

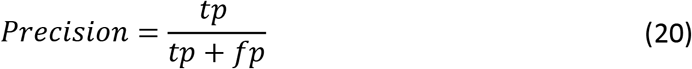

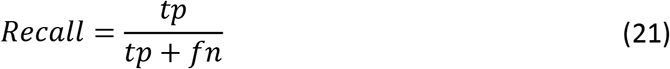

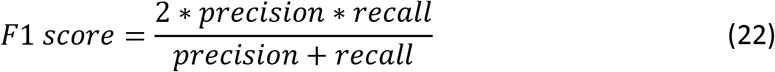

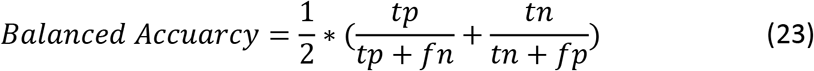

For any position of Env, we distinguish between two states: **(i)** Variance-positive (variance in amino acid sequence at the position is greater than 0), and **(ii)** No-variance (all sequences from the patient have the same amino acid at the position). Accordingly, our definitions are:

*tp*: Number of patients with a variance-positive position that are predicted as variance-positive.
*fp*: Number of patients with a no-variance position that are predicted as variance-positive.
*tn*: Number of patients with a no-variance position that are predicted as no-variance.
*fn*: Number of patients with a variance-positive position that are predicted as no-variance.

## 3. RESULTS

In this study, we tested the algorithm with patient-derived datasets from two HIV-1 subtypes. Specifically, we examined the ability of the algorithm to predict the level of variance at any position of Env based on the variance at the 10 closest positions on the three-dimensional structure of the protein. Initially, we tested performance for individual positions. We then proceeded to test performance for multi-position targets that represent the “footprint” of therapeutics on the Env protein.

We first compared the results of the KBC algorithm with those of the base learner (decision tree) and static ensemble (random forest). To increase the stringency of the comparison, we tuned both the decision tree and random forest algorithms for optimal performance. The shared hyperparameters between these models are the maximum depth of the trees and the minimum samples in the leaf nodes. Besides, we considered splitting criteria for the decision tree and the number of estimators for the random forest as the other hyperparameters for tuning. The grid search strategy was used to tune these models. Finally, we compared the performance of KBC with other classification techniques.

To estimate the evaluation metrics for each machine learning technique and each dataset, we used nested cross-validation to tune the hyperparameters and to obtain estimates of the classification metrics. We used k=5 for both the inner and outer folds.

### 3.1. Prediction of variance patterns in HIV-1 Env

New forms of the Env protein are continuously generated in HIV-infected individuals by the error-prone replication machinery of this virus. The appearance of new amino acid variants at Env positions targeted by therapeutics can lead to virus resistance to their effects. Such events appear to be random and thus considered unpredictable. There is a clinical need to understand the spatiotemporal patterns of variance in the HIV-infected host, which may lead to the development of new treatment strategies. We hypothesized that at any time in the infected host, positions that exhibit variance in amino acid sequence are spatially clustered on the Env protein. Such patterns are intuitive since the immune and fitness pressures mostly act on multi-position domains of Env rather than individual positions. Toward a better understanding of such patterns, we sought to determine whether the variance at any Env position can be accurately estimated based on the variance at adjacent positions on the protein.

To this end, we applied the KBC algorithm with sequences of Envs cloned from patient blood samples (6-30 Envs sequences were available and used for each sample). All sequences from each patient sample were aligned and compared to determine the absence or presence of in-host variance at each of the 856 positions of Env. As detailed below, we focused our analyses on Env positions targeted by therapeutics. The response variable is thus the absence or presence of variance at specific key positions of Env. The explanatory variable is the variance at the 10 positions closest to the key position on the protein (determined by the physical distance between the closest atoms of the two positions, measured in Ångtroms). The goal is to correctly classify the variance of a position using the variance of the adjacent positions. We decided to use the 10 adjacent positions since this is approximately the maximal number of amino acids that can contact the position of interest on the three-dimensional structure of Env. We note that the actual number of residues that are in contact or adjacent to each position may vary according to the location on the protein. For example, for any position buried within the core of the protein, its 10 nearest positions will be closer than for a position located on a loop that is exposed to the solvent. Nevertheless, we decided that as a first step, we will maintain this variable constant for all positions.

We first tested the ability of the KBC algorithm to predict the absence or presence of variance at individual positions in the high-mannose patch of Env (**Figure 4**). These N-linked glycans help to shield Env from recognition by host antibodies [49]; however, they also serve as targets for microbicidal agents such as lectins [50, 51] and therapeutic antibodies [52, 53]. We tested three positions in the high-mannose patch, namely positions 289, 332, and 339. These positions form part of the target sites for multiple agents that inhibit HIV-1, including antibodies 2G12, 10-1074, PGT135, PGT128, and DH270.5 [54–58], and the lectin microbicide griffithsin [59]. Data are composed of 1,960 amino acid sequences from 109 patients infected by HIV-1 subtype C, which is the most prevalent HIV-1 clade worldwide [60]. For position 289, the ratio of the variance-positive class to the no-variance class was 34:75. This ratio for positions 332 and 339 was 46:63 and 53:56, respectively.

**Figure 4.**
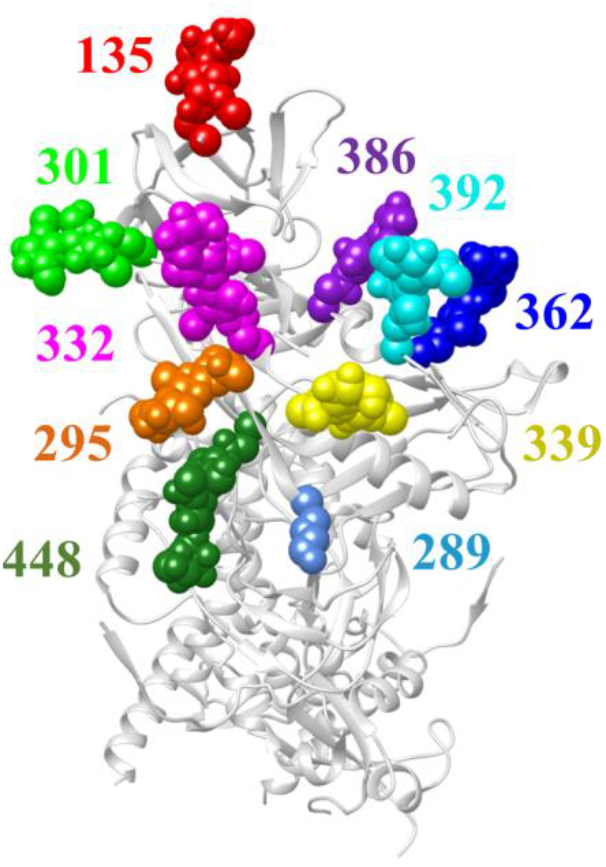
Cryo-EM structure of HIV-1 Env (side view, PDB ID 5FUU). Positions in the high-mannose patch are shown as spheres and labeled by Env position number. All positions shown contain N-linked glycans, except position 289, which contains Arg in the Env of HIV-1 isolate JRFL used to generate this structure.

For positions 289 and 339, the results of the KBC analyses showed improvement relative to the base learner (decision tree) and random forest (**Figure 5)**. By contrast, the prediction of variance at position 332 by the KBC method was similar to that of the other methods.

**Figure 5.**
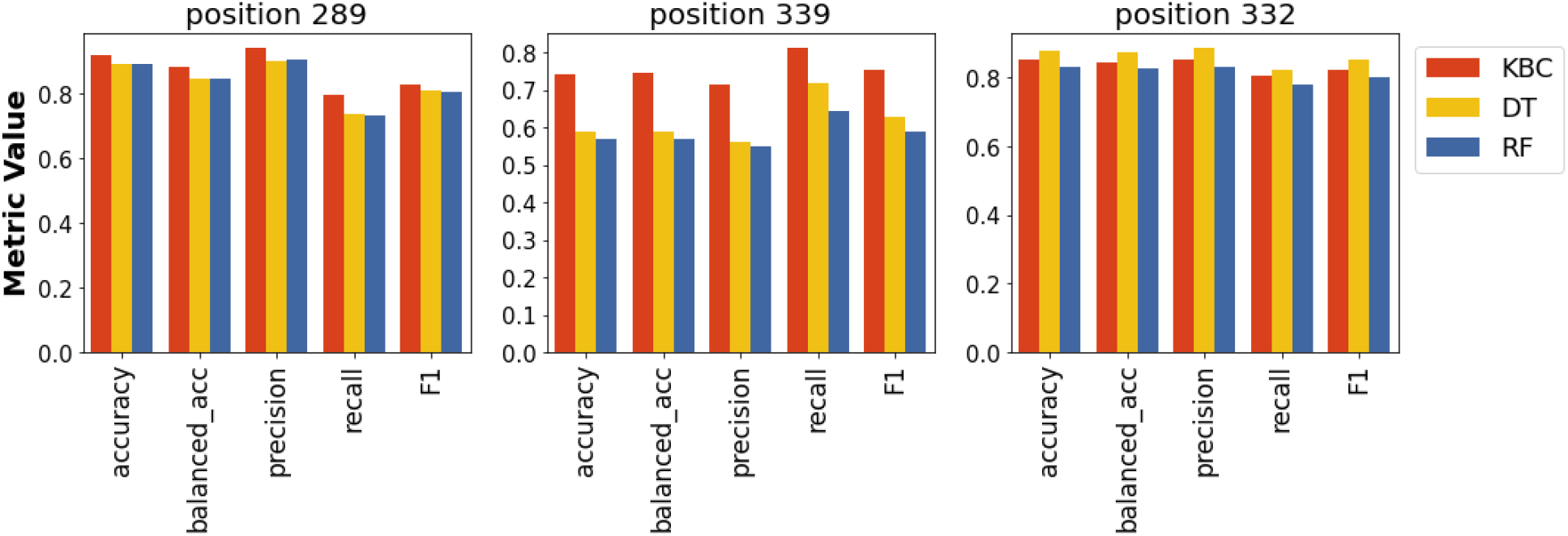
Predictions of variance at positions 289, 339, and 332 of Env using data from patients infected by HIV-1 clade C.

We also compared the performance of KBC with other machine learning algorithms (**Table 2**). Again, we observed modestly better performance of KBC for positions 289 and 339, whereas for position 332, the performance was similar to (or slightly worse than) other methods. We note that although KBC generally exhibited better point estimates than other methods, it also exhibited relatively high variability (see values in parentheses in **Table 2**).

**Table 2.** Prediction of variance at Env positions in the high-mannose patch by KBC and other algorithms.

| Position | Method <sup>a</sup> | Balanced Accuracy <sup>b,c</sup> | Accuracy | Precision | Recall | F1 Score |
| --- | --- | --- | --- | --- | --- | --- |
| 289 | KBC | **0.88** ($\pm 0.13$) | **0.92** ($\pm 0.08$) | **0.94** ($\pm 0.07$) | **0.80** ($\pm 0.26$) | 0.83 ($\pm 0.20$) |
| | QDA | 0.83 ($\pm 0.01$) | 0.88 ($\pm 0.01$) | 0.90 ($\pm 0.07$) | 0.71 ($\pm 0.05$) | 0.79 ($\pm 0.01$) |
| | LDA | 0.87 ($\pm 0.02$) | 0.91 ($\pm 0.01$) | 0.93 ($\pm 0.05$) | 0.76 ($\pm 0.05$) | **0.84** ($\pm 0.03$) |
| | NB | 0.71 ($\pm 0.18$) | 0.69 ($\pm 0.26$) | 0.66 ($\pm 0.26$) | 0.76 ($\pm 0.05$) | 0.67 ($\pm 0.16$) |
| | ADA | 0.86 ($\pm 0.01$) | 0.90 ($\pm 0.01$) | 0.91 ($\pm 0.07$) | 0.76 ($\pm 0.05$) | 0.83 ($\pm 0.02$) |
| | LogReg | 0.86 ($\pm 0.01$) | 0.90 ($\pm 0.01$) | 0.91 ($\pm 0.07$) | 0.76 ($\pm 0.05$) | 0.83 ($\pm 0.02$) |
| | SVM | 0.83 ($\pm 0.01$) | 0.88 ($\pm 0.01$) | 0.90 ($\pm 0.07$) | 0.71 ($\pm 0.05$) | 0.79 ($\pm 0.01$) |
| 339 | KBC | **0.75** ($\pm 0.12$) | **0.74** ($\pm 0.12$) | 0.71 ($\pm 0.12$) | 0.81 ($\pm 0.13$) | **0.75** ($\pm 0.11$) |
| | QDA | 0.59 ($\pm 0.07$) | 0.59 ($\pm 0.07$) | 0.56 ($\pm 0.09$) | 0.72 ($\pm 0.09$) | 0.63 ($\pm 0.07$) |
| | LDA | 0.57 ($\pm 0.03$) | 0.57 ($\pm 0.03$) | 0.55 ($\pm 0.05$) | 0.64 ($\pm 0.13$) | 0.59 ($\pm 0.06$) |
| | NB | 0.65 ($\pm 0.08$) | 0.65 ($\pm 0.07$) | 0.64 ($\pm 0.00$) | 0.68 ($\pm 0.16$) | 0.66 ($\pm 0.19$) |
| | ADA | 0.60 ($\pm 0.02$) | 0.60 ($\pm 0.02$) | 0.59 ($\pm 0.03$) | 0.62 ($\pm 0.06$) | 0.59 ($\pm 0.02$) |
| | LogReg | 0.59 ($\pm 0.06$) | 0.58 ($\pm 0.07$) | 0.54 ($\pm 0.05$) | **0.94** ($\pm 0.05$) | 0.69 ($\pm 0.03$) |
| | SVM | 0.65 ($\pm 0.04$) | 0.66 ($\pm 0.05$) | **1.00** ($\pm 0.09$) | 0.30 ($\pm 0.04$) | 0.48 ($\pm 0.02$) |
| 332 | KBC | 0.85 ($\pm 0.07$) | 0.85 ($\pm 0.07$) | 0.86 ($\pm 0.12$) | 0.80 ($\pm 0.08$) | 0.83 ($\pm 0.08$) |
| | QDA | 0.84 ($\pm 0.05$) | 0.84 ($\pm 0.06$) | 0.84 ($\pm 0.12$) | 0.83 ($\pm 0.11$) | 0.82 ($\pm 0.06$) |
| | LDA | 0.87 ($\pm 0.03$) | 0.88 ($\pm 0.03$) | 0.89 ($\pm 0.05$) | 0.83 ($\pm 0.06$) | 0.85 ($\pm 0.04$) |
| | NB | 0.61 ($\pm 0.16$) | 0.57 ($\pm 0.21$) | 0.59 ($\pm 0.23$) | **0.91** ($\pm 0.13$) | 0.67 ($\pm 0.10$) |
| | ADA | 0.87 ($\pm 0.03$) | 0.88 ($\pm 0.03$) | 0.89 ($\pm 0.05$) | 0.83 ($\pm 0.06$) | 0.85 ($\pm 0.04$) |
| | LogReg | 0.82 ($\pm 0.09$) | 0.81 ($\pm 0.12$) | 0.80 ($\pm 0.18$) | 0.87 ($\pm 0.11$) | 0.81 ($\pm 0.08$) |
| | SVM | **0.88** ($\pm 0.02$) | **0.89** ($\pm 0.02$) | **0.89** ($\pm 0.05$) | 0.85 ($\pm 0.03$) | **0.87** ($\pm 0.03$) |

<sup>a</sup> Calculations were performed using data from 109 patients infected by HIV-1 clade C.
<sup>b</sup> Standard deviation values are indicated in parentheses.
<sup>c</sup> Values in bold font indicate the highest point estimation value for each metric.

This likely occurred due to the relatively small size of the dataset. Later, we show that increasing the size of the dataset drastically reduces the variability of the estimates.

Antiviral therapeutics bind to targets composed of multiple residues; their “footprint” on the viral protein can span a large surface that contains multiple amino acids [61–64]. Changes at any of these contacts may reduce Env recognition by the therapeutic and cause resistance. We examined the performance of the KBC algorithm to predict variance in a combined feature composed of 10 positions in the high-mannose patch shown in **Figure 4**. To this end, for each position *A_i_* in the high-mannose patch (*i* = 1,2,…, 10), we relabeled its 10 adjacent positions as 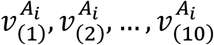, where 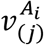 is the variance at the *j-th* adjacent position to *A_i_* (*j* = 1,2,…, 10). For each *j*, we then combined the 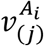 values of the 10 *A_i_* positions.

We first used the dataset of sequences from 109 HIV-1 clade C infected individuals. Results were compared between the KBC method and the above machine learning methods. The ratio of positive-variance to the no-variance instances for this dataset was 450:640. Remarkably, KBC performed better than all models to predict sequence variance in the high-mannose patch **(Table 3)**.

**Table 3.** Prediction of variance in the high-mannose patch of Env by KBC and other algorithms.

| Clade | Method | Balanced Accuracy<sup>a,b</sup> | Accuracy | Precision | Recall | F1 Score |
| --- | --- | --- | --- | --- | --- | --- |
| Clade C | KBC | **0.65** ($\pm 0.05$) | **0.69** ($\pm 0.05$) | **0.67** ($\pm 0.08$) | 0.47 ($\pm 0.05$) | 0.55 ($\pm 0.08$) |
| | DT | 0.59 ($\pm 0.05$) | 0.63 ($\pm 0.07$) | 0.54 ($\pm 0.07$) | 0.39 ($\pm 0.21$) | 0.43 ($\pm 0.17$) |
| | RF | 0.59 ($\pm 0.02$) | 0.63 ($\pm 0.03$) | 0.60 ($\pm 0.05$) | 0.34 ($\pm 0.13$) | 0.42 ($\pm 0.10$) |
| | QDA | 0.60 ($\pm 0.02$) | 0.63 ($\pm 0.04$) | 0.57 ($\pm 0.02$) | 0.39 ($\pm 0.16$) | 0.45 ($\pm 0.12$) |
| | LDA | 0.58 ($\pm 0.04$) | 0.62 ($\pm 0.06$) | 0.54 ($\pm 0.07$) | 0.39 ($\pm 0.17$) | 0.44 ($\pm 0.13$) |
| | NB | 0.61 ($\pm 0.06$) | 0.63 ($\pm 0.08$) | 0.54 ($\pm 0.08$) | 0.52 ($\pm 0.21$) | 0.52 ($\pm 0.14$) |
| | ADA | 0.50 ($\pm 0.05$) | 0.44 ($\pm 0.04$) | 0.42 ($\pm 0.02$) | **0.88** ($\pm 0.09$) | **0.56** ($\pm 0.02$) |
| | LogReg | 0.57 ($\pm 0.06$) | 0.61 ($\pm 0.04$) | 0.55 ($\pm 0.07$) | 0.33 ($\pm 0.20$) | 0.38 ($\pm 0.15$) |
| | SVM | 0.58 ($\pm 0.04$) | 0.62 ($\pm 0.06$) | 0.56 ($\pm 0.06$) | 0.35 ($\pm 0.23$) | 0.40 ($\pm 0.17$) |
| Clade B | KBC | 0.65 ($\pm 0.02$) | **0.74** ($\pm 0.02$) | **0.68** ($\pm 0.06$) | 0.39 ($\pm 0.03$) | 0.50 ($\pm 0.04$) |
| | DT | 0.60 ($\pm 0.05$) | 0.70 ($\pm 0.02$) | 0.61 ($\pm 0.03$) | 0.29 ($\pm 0.16$) | 0.36 ($\pm 0.16$) |
| | RF | 0.62 ($\pm 0.00$) | 0.72 ($\pm 0.00$) | 0.61 ($\pm 0.02$) | 0.35 ($\pm 0.02$) | 0.45 ($\pm 0.01$) |
| | QDA | 0.63 ($\pm 0.01$) | 0.73 ($\pm 0.01$) | 0.64 ($\pm 0.03$) | 0.36 ($\pm 0.02$) | 0.46 ($\pm 0.02$) |
| | LDA | 0.64 ($\pm 0.01$) | 0.72 ($\pm 0.01$) | 0.61 ($\pm 0.01$) | 0.40 ($\pm 0.04$) | 0.48 ($\pm 0.03$) |
| | NB | **0.67** ($\pm 0.04$) | 0.70 ($\pm 0.03$) | 0.53 ($\pm 0.05$) | 0.59 ($\pm 0.07$) | **0.56** ($\pm 0.06$) |
| | ADA | 0.41 ($\pm 0.05$) | 0.29 ($\pm 0.02$) | 0.28 ($\pm 0.03$) | **0.74** ($\pm 0.14$) | 0.40 ($\pm 0.05$) |
| | LogReg | 0.64 ($\pm 0.02$) | 0.72 ($\pm 0.01$) | 0.61 ($\pm 0.02$) | 0.39 ($\pm 0.05$) | 0.47 ($\pm 0.03$) |
| | SVM | 0.63 ($\pm 0.02$) | 0.70 ($\pm 0.02$) | 0.55 ($\pm 0.03$) | 0.41 ($\pm 0.01$) | 0.47 ($\pm 0.02$) |

<sup>a</sup> Standard deviation values are indicated in parentheses.
<sup>b</sup> Values in bold font indicate the highest point estimation value for each metric.

To validate these results, we examined the ability of KBC to predict variance in a second independent panel of sequences derived from individuals infected by HIV-1 clade B. This clade is the most prevalent in the United States and Europe [60]. Sequences from 191 patients were tested to predict variance at the multi-position high-mannose patch using the different algorithms. Consistent with the data shown for clade C, the performance of KBC was superior, albeit modestly, to that of the other algorithms (**Table 3**). The ratio of the positive-variance class to the no-variance class for the clade B dataset was 621:1289.

We further expanded our studies to test a second clinically significant domain of the Env protein, namely the CD4-binding site. This domain interacts with the receptor for the virus, which allows entry of the viral genome into the cell [65]. Since this site is conserved among diverse HIV-1 strains, it also serves as a target for multiple therapeutics, including the small molecule Fostemsavir [38] and antibody therapeutics VRC01 and 3BNC117 [39, 40]. We tested a combination of the 23 positions that serve as the contact sites for both antibodies VRC01 and 3BNC117 (**Figure 6**). We applied the same procedure explained for the high-mannose patch positions to combine the positions of the CD4-binding site.

**Figure 6.**
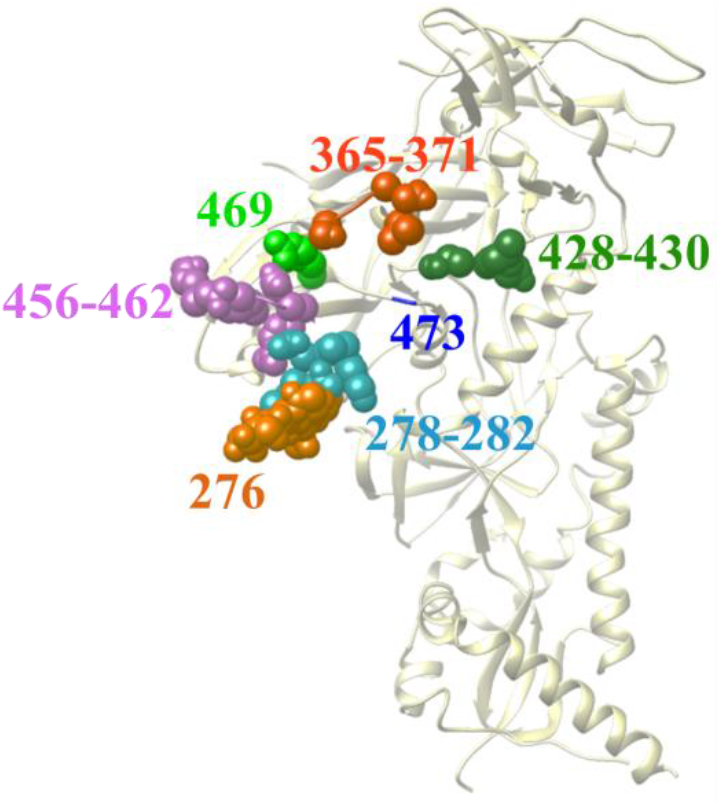
Side view of the cryo-EM structure of HIV-1 Env (PDB ID 5FUU). Positions in the CD4-binding site contacted by antibodies 3BNC117 and VRC01 are shown as spheres and labeled.

The ratio of positive-variance to the no-variance classes for the CD4-binding site dataset was 685:3708 and 557:1950 for clades B and C, respectively. The performance of KBC was compared with all other algorithms tested above. Interestingly, the performance of the KBC algorithm was considerably higher than that of other algorithms (**Table 4**). For positions in the CD4-binding site, the increase in performance was better than that observed for positions in the high-mannose patch (**Table 3**). Comparison of the results in **Table 3** and **Table 4** shows that the variability of the estimates was considerably lower when we analyzed a group of positions rather than individual positions. Thus, for the CD4-binding site, the standard deviation in accuracy, balanced accuracy, recall, and F1 score obtained by KBC is the smallest among all other models for clade C. Indeed, KBC shows higher point estimates as well as smaller standard deviation values for the estimates.

**Table 4.** Prediction of variance in the CD4-binding site of Env using KBC and other algorithms.

| Clade | Methods | Balanced Accuracy<sup>a,b</sup> | Accuracy | Precision | Recall | F1 Score |
| --- | --- | --- | --- | --- | --- | --- |
| Clade C | KBC | **0.71** ($\pm 0.01$) | **0.85** ($\pm 0.01$) | **0.81** ($\pm 0.07$) | 0.45 ($\pm 0.03$) | **0.58** ($\pm 0.02$) |
| | DT | 0.61 ($\pm 0.13$) | 0.76 ($\pm 0.03$) | 0.32 ($\pm 0.23$) | 0.34 ($\pm 0.42$) | 0.25 ($\pm 0.25$) |
| | RF | 0.56 ($\pm 0.04$) | 0.76 ($\pm 0.03$) | 0.49 ($\pm 0.08$) | 0.19 ($\pm 0.18$) | 0.22 ($\pm 0.14$) |
| | QDA | 0.59 ($\pm 0.07$) | 0.75 ($\pm 0.04$) | 0.50 ($\pm 0.08$) | 0.29 ($\pm 0.28$) | 0.28 ($\pm 0.15$) |
| | LDA | 0.69 ($\pm 0.11$) | 0.79 ($\pm 0.03$) | 0.59 ($\pm 0.09$) | 0.50 ($\pm 0.31$) | 0.46 ($\pm 0.19$) |
| | NB | 0.67 ($\pm 0.09$) | 0.73 ($\pm 0.09$) | 0.45 ($\pm 0.11$) | **0.58** ($\pm 0.31$) | 0.45 ($\pm 0.16$) |
| | ADA | 0.46 ($\pm 0.18$) | 0.62 ($\pm 0.24$) | 0.44 ($\pm 0.29$) | 0.15 ($\pm 0.10$) | 0.20 ($\pm 0.14$) |
| | LogReg | 0.65 ($\pm 0.12$) | 0.79 ($\pm 0.01$) | 0.56 ($\pm 0.04$) | 0.39 ($\pm 0.33$) | 0.38 ($\pm 0.21$) |
| | SVM | 0.64 ($\pm 0.11$) | 0.76 ($\pm 0.03$) | 0.50 ($\pm 0.06$) | 0.43 ($\pm 0.35$) | 0.37 ($\pm 0.16$) |
| Clade B | KBC | **0.69** ($\pm 0.02$) | **0.89** ($\pm 0.01$) | **0.78** ($\pm 0.02$) | 0.40 ($\pm 0.04$) | **0.53** ($\pm 0.04$) |
| | DT | 0.54 ($\pm 0.02$) | 0.82 ($\pm 0.02$) | 0.33 ($\pm 0.02$) | 0.13 ($\pm 0.08$) | 0.17 ($\pm 0.10$) |
| | RF | 0.53 ($\pm 0.02$) | 0.82 ($\pm 0.03$) | 0.35 ($\pm 0.18$) | 0.11 ($\pm 0.05$) | 0.15 ($\pm 0.06$) |
| | QDA | 0.56 ($\pm 0.03$) | 0.82 ($\pm 0.04$) | 0.42 ($\pm 0.16$) | 0.17 ($\pm 0.11$) | 0.21 ($\pm 0.09$) |
| | LDA | 0.58 ($\pm 0.05$) | 0.82 ($\pm 0.01$) | 0.38 ($\pm 0.05$) | 0.24 ($\pm 0.13$) | 0.28 ($\pm 0.10$) |
| | NB | 0.64 ($\pm 0.08$) | 0.75 ($\pm 0.10$) | 0.34 ($\pm 0.14$) | 0.47 ($\pm 0.21$) | 0.37 ($\pm 0.13$) |
| | ADA | 0.48 ($\pm 0.03$) | 0.17 ($\pm 0.02$) | 0.15 ($\pm 0.01$) | **0.93** ($\pm 0.10$) | 0.26 ($\pm 0.02$) |
| | LogReg | 0.56 ($\pm 0.04$) | 0.84 ($\pm 0.00$) | 0.42 ($\pm 0.05$) | 0.17 ($\pm 0.10$) | 0.23 ($\pm 0.11$) |
| | SVM | 0.55 ($\pm 0.04$) | 0.82 ($\pm 0.03$) | 0.35 ($\pm 0.15$) | 0.17 ($\pm 0.11$) | 0.21 ($\pm 0.10$) |

<sup>a</sup> Standard deviation values are indicated in parentheses.
<sup>b</sup> Values in bold font indicate the highest point estimation value for each metric.

Taken together, these findings show that when Env positions are tested individually, KBC outperforms other algorithms for most (but not all) positions. Nevertheless, this algorithm shines in its performance when tested with a combination of positions that describe the complex (multi-position) target sites of the therapeutics on the Env protein.

### 3.2. KBC hyperparameter analysis

We examined the effects of two important hyperparameters of the KBC model on its performance. Data that describe variance patterns in the high-mannose patch were used. To evaluate the performance, we used the balanced accuracy metric. We explored the effect of one hyperparameter while maintaining the rest at a constant level. We used 20 decision trees (M=20); for each, we picked 4 features randomly (*f*=4), and the maximum depth was set to be 4. The OOB weight was fixed for both experiments at its default value, 0.632.

First, we explored the effect of the minimum acceptance threshold (*δ*). For this experiment, the number of neighbors was set to 10. The experiment was conducted with a variety of thresholds from 0 to 1, and the balanced accuracy was calculated. We observed that for the clade B dataset, increasing the minimum acceptance threshold improved the performance of the KBC model (**Figure 7A**). For the clade C dataset, the performance also increased gradually; however, it peaked at a threshold of 0.65, followed by a modest reduction (**Figure 7B**). These findings suggest that increasing the value for *δ* results in an overall increase in performance due to the higher confidence in the set of selected classifiers. However, in some cases, further increases in *δ* may result in loss of useful learners that can reduce overall performance.

**Figure 7.**
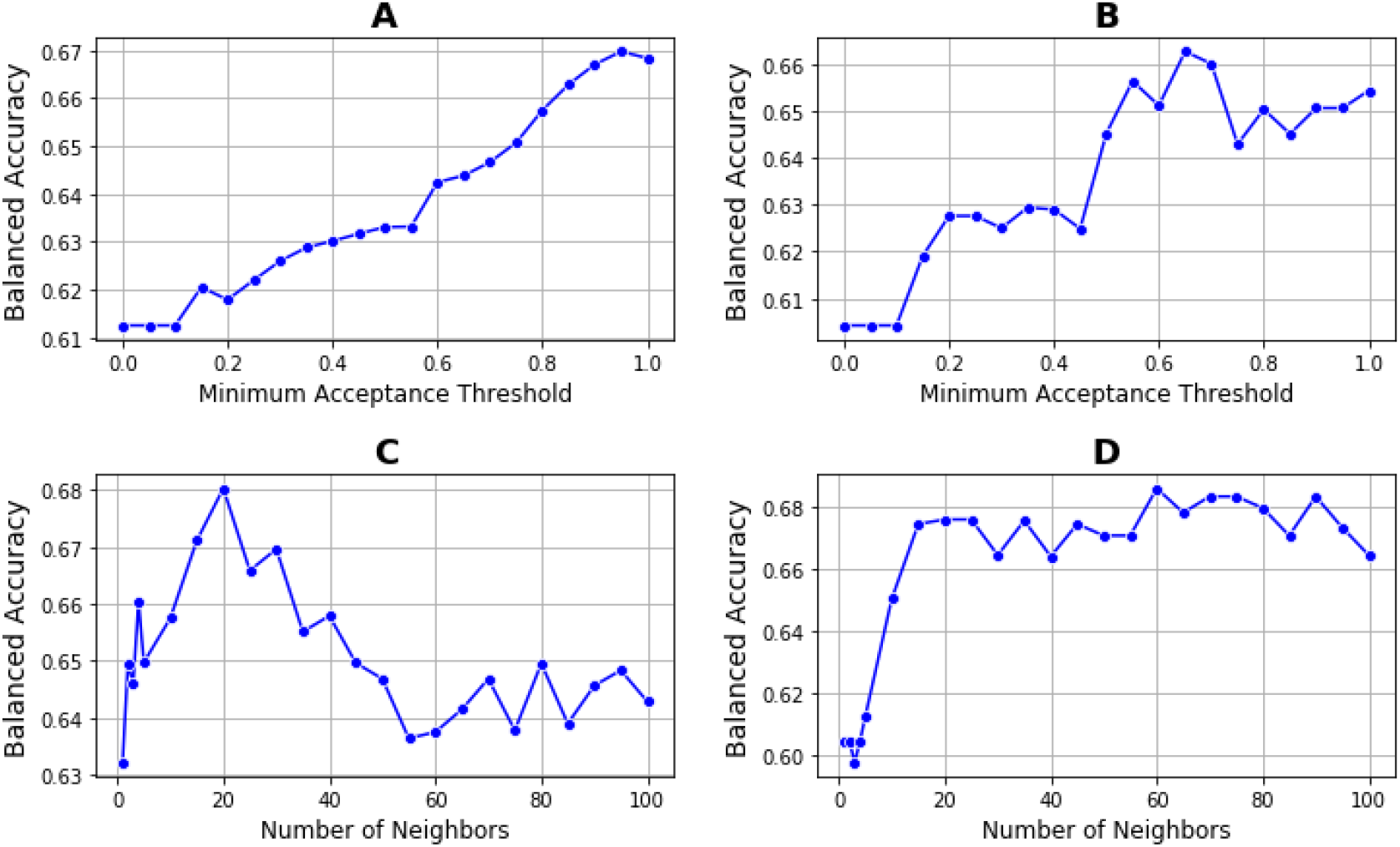
(A) Effect of the minimum acceptance threshold on the balanced accuracy using data that describe variance patterns in the high-mannose patch in clade B (B) Effect of the minimum acceptance threshold on the balanced accuracy using data that describe variance patterns in the high-mannose patch in clade C, (C) Effect of the neighborhood size on the balanced accuracy using data that describe variance patterns in the CD4-binding site in clade B, and (D) Effect of the neighborhood size on the balanced accuracy using data that describe variance patterns in the CD4-binding site in clade C.

We also explored the effect of the neighborhood size on the performance of the KBC algorithm. Here we used *δ*=0.8 as the minimum acceptance threshold. Different numbers of neighbors (ranging between 1 and 100) were tested to examine the effect of neighborhood size on the performance of the model. We observed that for both clades B and C, increasing the number of neighbors up to approximately 15 or 20 increased the performance (**Figure 7, C,** and **D**). Further increases in the neighborhood size decreased the performance in clade B, whereas it did not impact clade C. These findings suggested that a neighborhood size of approximately 15 is optimal for the data that describe variance patterns in high-mannose patch given the above hyperparameters.

### 3.3. Effect of base learners on performance of the KBC algorithm

As a further analysis, we examined whether the choice of base learner in the KBC algorithm affects the overall performance of the method. To this end, we used logistic regression and Naïve Bayes (separately) as the base learners. We evaluated the KBC method using data from HIV-1 clade C that describe variance patterns in the high-mannose patch and the CD4-binding site. These results were compared with the results obtained using decision tree as the base learner. In this experiment, we kept the structure of the KBC algorithm as before, with the exception that for each trial a homogenous set of base learners from one type was utilized (i.e., decision tree, logistic regression, or Naïve Bayes). For the tuning process and for the KBC with logistic regression as the base learner, we incorporated the hyperparameter *C*, which is the inverse of the regularization strength. For the KBC method with Naïve Bayes as the base learner, no hyperparameter was added to the KBC’s hyperparameter list.

Results of the above tests are shown in **Figure 8**. For the high-mannose patch (**Figure 8A**), decision tree yielded modestly higher point estimates for accuracy and precision, whereas Naïve Bayes showed modestly better recall and F1 score. However, these differences were not statistically significant (see error bars in **Figure 8**). Therefore, for this dataset, the choice of base learner did not impact the performance of the KBC method. For the CD4-binding site (**Figure 8B**), decision tree and logistic regression performed equally well as the base learners and were both better than Naïve Bayes in accuracy and precision metrics. Similar to the high-mannose-patch data, the recall was better for Naïve Bayes; however, this improvement was not sufficient to counterbalance the considerably lower precision, resulting in an F1 score for Naïve Bayes that was modestly smaller than that of decision tree and logistic regression.

**Figure 8.**
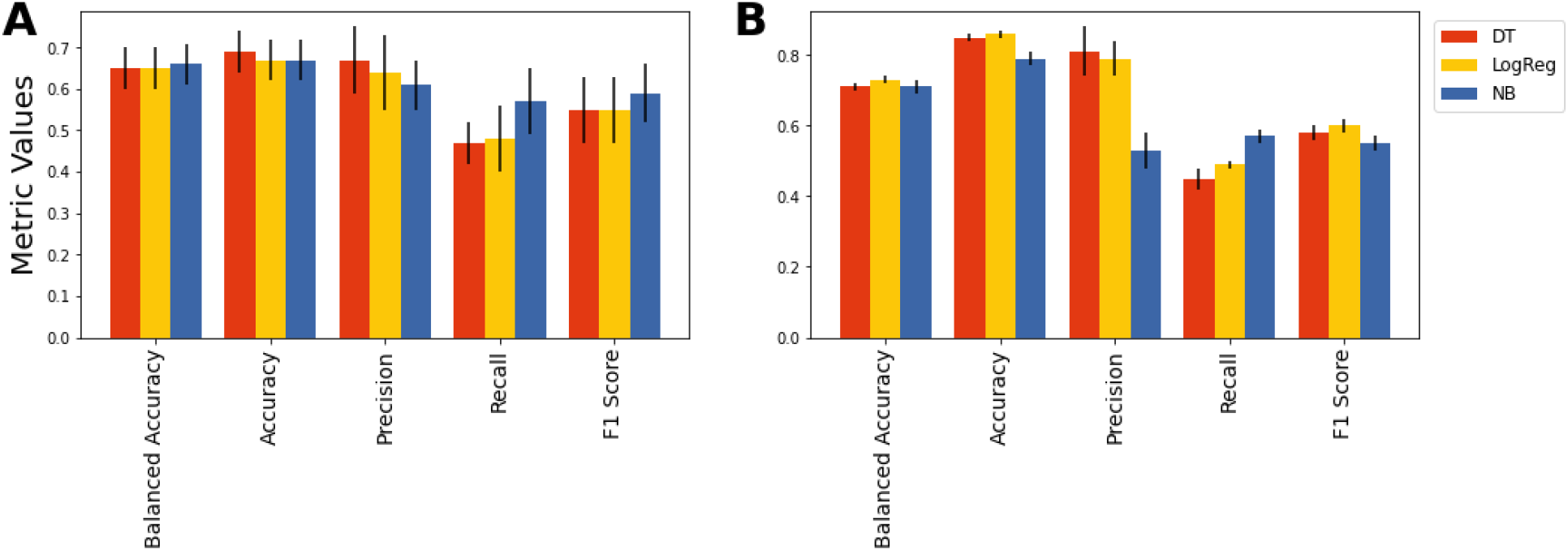
Performance of the KBC algorithm using decision tree, logistic regression, and Naïve Bayes as the base learners with data from HIV-1 clade C that describe variance patterns in the high-mannose patch (A) and the CD4-binding site (B). Error bars indicate standard deviation.

In summary, KBC is a general framework in which any choice of base learners can be used. As shown in **Figure 8**, the choice of base learner may affect the performance of the KBC algorithm. These effects are likely specific for each problem. In this study, we utilized decision tree as the base learner due to its speed and performance that was at least as good as other options.

## 4. DISCUSSION

Many viruses, including HIV-1, exhibit a high error rate during their replication [66, 67]. New variants of their proteins are continuously generated in the host. The ability to create diversity allows viruses to rapidly adapt to selective pressures, including antiviral therapeutics. The first step in the emergence of resistance is the appearance of sequence variance at a position of the viral protein targeted by the therapeutic. Variance patterns across the Env protein seem random and are thus considered unpredictable. In these studies, we examined whether positions that exhibit sequence variance are spatially clustered on the three-dimensional structure of the HIV-1 Env protein. Specifically, we tested whether the absence or presence of sequence variance at any position of Env in a patient can be predicted by variance at adjacent positions on the protein. To address this question, we developed a new dynamic ensemble selection algorithm based on bootstrap resampling.

We used the KBC method, which defines the neighborhood of a new data point using the KNN algorithm. Specifically, for each position of interest, KBC defines the neighborhood by identifying observations that have similar feature vectors (i.e., a similar variance profile of the 10 adjacent positions). We then selected the k-best classifier(s) within that neighborhood based on a weighted score procedure. By comparing each classifier’s score with a minimum acceptance threshold, we obtain the set of best classifiers to predict the class label for each new instance. The dynamism, along with the specific design, resulted in a flexible approach that is not constrained to select a constant number of learners every time it wants to classify a new observation. Therefore, based on the performance of the learners, only those classifiers surpassing an explicit expectation are chosen; this results in an improvement in the overall performance. The novelty of this algorithm is in the dynamic classifier selection mechanism, in which we designed a weighting procedure to evaluate each classifier’s performance within a neighborhood of an instance and decide if the classifier is good enough to classify the observation. This approach is based on bootstrap resampling, which creates out-of-bag samples that can be used along with the resampled data in the classifiers’ evaluation process.

We applied the algorithms to predict the level of variance at individual positions of Env based on variance at adjacent positions on the molecule. Results were compared with a variety of state-of-art methods, such as the Adaboost, Naïve Bayes, logistic regression, linear and quadratic discriminant analysis methods, and SVM. Overall, the KBC algorithm predicted the absence or presence of variance better than the above machine learning tools. Interestingly, performance varied with the domain of Env tested. Only modest enhancement of performance by the KBC method was observed for the high-mannose patch of Env, whereas dramatic enhancement was observed for the CD4-binding site. Improvement in all classification metrics was observed.

Importantly, KBC showed considerable improvement for predicting variance at multi-position features. We tested two Env domains targeted by therapeutics; the CD4-binding site and the high-mannose patch of Env (composed of 23 and 10 positions, respectively). Both domains constitute targets for multiple HIV-1 therapeutics [39, 40, 54–58]. These antibody footprints on Env were analyzed using sequence data from patients infected by HIV-1 clades B and C, which were analyzed separately. For both domains and in both clades, the absence or presence of variance was predicted better using KBC than other algorithms. These results are highly encouraging since therapeutics do not recognize single positions but rather multi-position footprints on the protein; a change at any position can reduce binding of the agents and increase clinical resistance [61, 62, 68]. The ability to predict the variance in a given domain based on the adjacent sites suggests that if these associations are stable over time, they may provide insight into future changes that may occur based on the current patterns of variance in the patient. Interestingly, for small datasets (e.g., analysis of single Env positions), KBC exhibited high point estimates but also high variability. By contrast, using larger datasets (e.g., multi-position targets), KBC exhibited both higher estimates and also smaller variability compared to the other algorithms. This finding suggested that the KBC model performs better with large datasets.

We observed that despite using homogenous and simple learners, KBC competes well with even sophisticated algorithms such as SVM, Adaboost, and discriminant analysis techniques. We also evaluated the effects of using logistic regression and Naïve Bayes as the base learners. Differences in the performance of KBC with each of these base learners were explored. Our results suggested that the choice of base learner may impact the overall performance; however, the effects are likely specific for each problem. We selected to focus most of our studies on decision tree as the base learner because of its relative speed and its performance, which was at least as high as that of the other options. Nevertheless, we note that by using more advanced methods as the base learner and by increasing diversity using a pool of different methods, KBC may exhibit even higher performance, which can be explored in future studies.

It should be noted that this study is subjected to a few limitations which suggest future research directions. First, the entire training dataset needs to be scanned for every new instance to find the neighbors with KNN. This may lead to computational intractability for very large datasets. Innovative methods for defining the neighborhood can be studied to improve efficiency. An example of such methods could be clustering algorithms that group similar instances into the same clusters [31]. Then, for each new observation, one can find the most similar cluster to the new data point and evaluate the classifiers within that neighborhood. Second, although the algorithm is capable of accommodating various configurations for the base learners, adding a high number of sophisticated methods can result in an increase in the training time for the model. Therefore, optimizing the choices for base learners would be necessary to balance running time with classification performance. Finally, we observed that for small datasets, KBC shows higher variability for the estimates compared to the other machine learning techniques. To address this issue, one may incorporate variance reduction techniques in the algorithm to decrease the variability for an estimate of interest such that it can be used for datasets with few observations.

